# Mice colonized with the defined microbial community OMM19.1 are susceptible to *C. difficile* infection without prior antibiotic treatment

**DOI:** 10.1101/2024.08.27.609948

**Authors:** Michelle Chua, James Collins

## Abstract

Diverse gut microorganisms present in humans and mice are essential for the prevention of microbial pathogen colonization. However, antibiotic-induced dysbiosis of the gut microbiome reduces microbial diversity and allows *C. difficile* to colonize the intestine. The Oligo Mouse Microbiota 19.1 (OMM19.1) is a synthetic community that consists of bacteria that are taxonomically and functionally designed to mimic the specific pathogen-free (SPF) mouse gut microbiota. Here, we examined the susceptibility of OMM19.1 colonized mice to *C. difficile* infection at a range of infectious doses (10^3^, 10^5^, and 10^7^ spores) without prior antibiotic treatment. We found that mice colonized with OMM19.1 were susceptible to *C. difficile* infection regardless of the dose. The clinical scores increased with increasing *C. difficile* dosage. Infection with *C. difficile* was correlated with a significant increase in *Ligilactobacillus murinus* and *Escherichia coli*, while the abundance of *Bacteroides caecimuris, Akkermansia muciniphila, Extibacter muris, and Turicimonas muris* significantly decreased following *C. difficile* infection. Our results demonstrate that the OMM19.1 community requires additional bacteria to enable colonization resistance.

## IMPORTANCE

The human gut microbiota consists of a wide range of microorganisms whose composition and function vary according to their location and have a significant impact on health and disease. The ability to generate and test defined microbiota within gnotobiotic animal models is essential for determining the mechanisms responsible for colonization resistance. The exact mechanism(s) by which healthy microbiota prevents *C. difficile* infection (CDI) is unknown, although competition for nutrients, active antagonism, production of inhibitory metabolites (such as secondary bile acids), and microbial manipulation of the immune system are all thought to play a role. Here, we colonized germ-free C57BL/6 mice with a synthetic bacterial community (OMM19.1) that mimics the SPF mouse microbiota. Following breeding, to enable immune system development, F1 mice were infected with multiple doses of *C. difficile*. Our research suggests that there are additional essential microbial functions that are absent from the current OMM19.1 model.

## OBSERVATION

The Oligo-Mouse-Microbiota 19.1 (OMM19.1) synthetic community builds upon the widely used Oligo-Mouse-Microbiota 12 (OMM12) (1), with the addition of nine species (2) (Table 1). This new 21 member community more accurately reflects SPF mouse microbiota, accounting for the absence of important microbial functions, including 7α-dehydroxylation via the addition of *Extibacter muris* (2, 3) and compensation for phenotypic differences between OMM12 and SPF mice, including body composition and immune cells in the intestine and associated lymphoid tissues (1). While the original OMM12 community provided colonization resistance against *Salmonella enterica* serovar Typhimurium infection (1), only partial resistance to *C. difficile* was observed after the addition of *Clostridium scindens* (4). *C. scindens* can modify primary bile acids (a key *C. difficile* germination trigger) via 7α-dehydroxylation and produces other bile acid-independent mechanisms that inhibit *C. difficile* outgrowth (5). Here, we examined the resistance of OMM19.1 to *C. difficile* infection (CDI).

**Table 1.** Strains in OMM19.1.

| Strain | DSM number |
| --- | --- |
| *Akkermansia muciniphila* | 26127 |
| *Bacteroides caecimuris* | 26085 |
| *Blautia pseudococcoides* | 26115 |
| *Enterocloster clos-*<br>*tridioformis* | 26114 |
| *Clostridium innocuum* | 26113 |
| *Enterococcus faecalis* | 32036 |
| *Flavonifractor plautii* | 26117 |
| *Limosilactobacillus reuteri* | 23035 |
| *Muribaculum intestinale* | 28989 |
| *Turicimonas muris* | 26109 |
| ***Acutalibacter muris*** | 26090 |
| ***Bifidobacterium animalis*** | 26074 |
| ***Thomasclavelia ramosa*** | 29357 |
| ***Adlercreutzia mucosicola*** | 19490 |
| ***Escherichia coli*** | 28618 |
| ***Extibacter muris*** | 28560 |
| ***Flintibacter butyricus*** | 27579 |
| ***Ligilactobacillus murinus*** | 28683 |
| ***Mucispirillum schaedleri*** | 104751 |
| ***Parabacteroides goldsteinii*** | 29187 |
| ***Xylanibacter rodentium*** | 105243 |

Germ-free C57BL/6 mice were obtained from the Functional Microbiomics, Inflammation, and Pathogenicity Center (University of Louisville) and were stably colonized with the 21 strain OMM19.1. After breeding, the F1 generation of OMM19.1 mice were used for *C. difficile* infection experiments.

Mice received low (10^3^ cfu), medium (10^5^ cfu), or high (10^7^ cfu) doses of *C. difficile* CD2015 (RT027) spores via oral gavage, without prior antibiotic treatment. All mice were susceptible to CDI regardless of the dose, with weight loss peaking on days 2, 3, and 4 post-infection in mice receiving low, medium, and high doses, respectively (Figure 1A). Clinical scores correlated with the *C. difficile* inoculum, with mice receiving the high dose exhibiting significantly higher disease scores than the low dose mice on day 1 (median [IQR], 1.5 [1] vs. 3 [0.5], p = 0.026). Mice that received the low inoculum recovered quicker. Mice receiving a medium- or high-dose had significantly higher disease scores than low-dose mice on day 4 (median [IQR], 2 [2.5] vs. 6 [1], p = 0.011, and 2 [2.5] vs. 6 [1.5], p = 0.008, Figure 1B). On day 1 post-infection, *C. difficile* levels were significantly different between groups (p = 0.002), with 2.0 × 10^5^ (SD 2.7 × 10^5^), 7.2 × 10^5^ (SD 6.9 × 10^5^), and 7.9 × 10^6^ (SD 5.5 × 10^6^) CFU/g *C. difficile* in the low, medium, and high dose groups respectively. No significant difference in *C. difficile* burden was observed at later timepoints (Figure 1C). No mice succumbed to the disease or met the euthanasia criteria.

**Figure 1.**
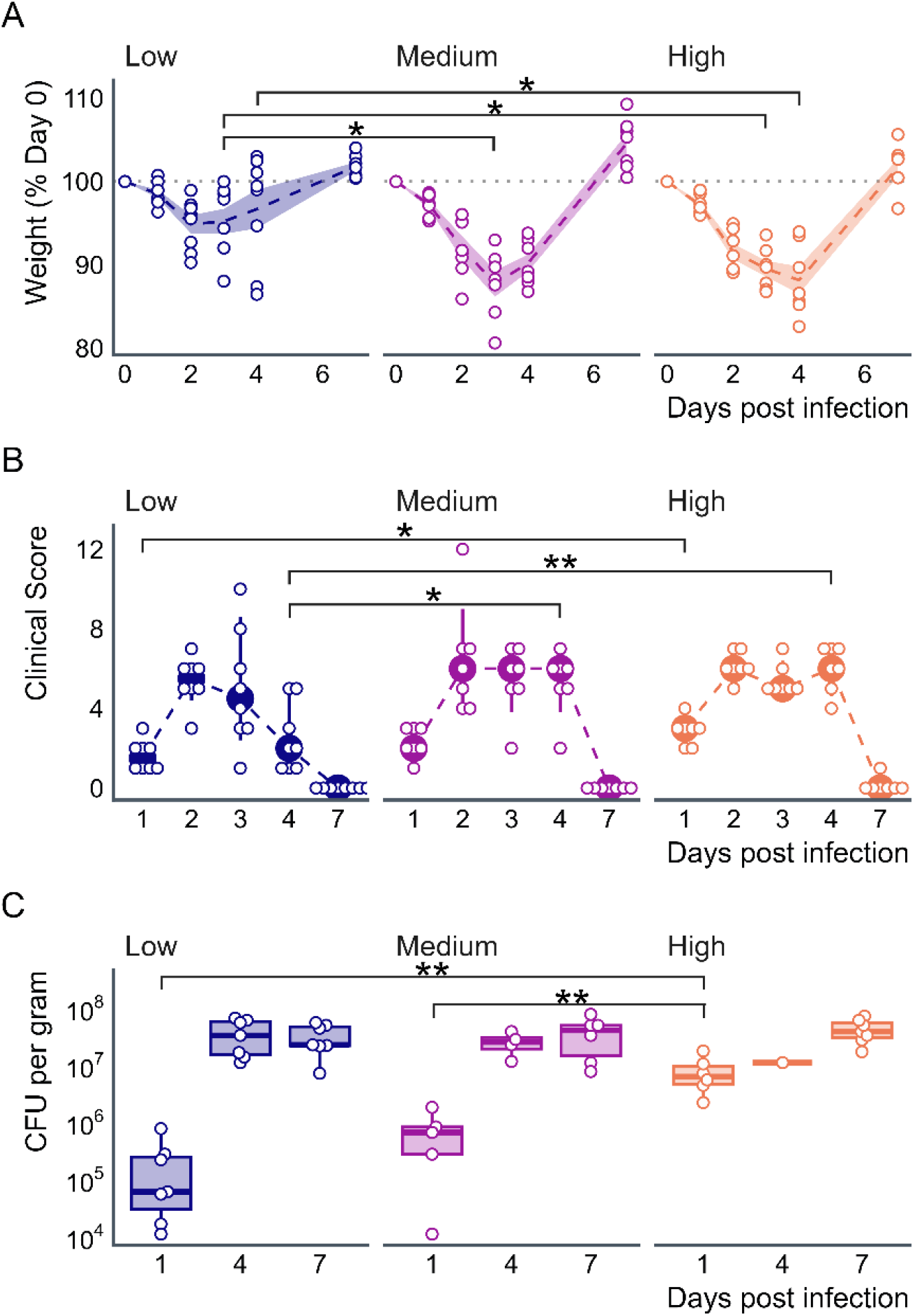
Effect of *C. difficile* inoculation dose on OMM19.1 mice. OMM19.1 mice were challenged with low (10^3^ cfu, n = 8), medium (10^5^ cfu, n = 7), or high (10^7^ cfu, n = 7) doses of *C. difficile*. **A) Weight loss**: Peak weight loss occurred on days 2, 3, and 4 in the low-, medium-, and high-dose groups, respectively, with average body weight losses of 5.1% (SD = 3.1), 12.3% (SD = 4.1), and 11.8% (SD = 4.3). Significant differences in weight were observed on day 3 post-infection (PI) between the low and medium groups (p = 0.028) and the low and high groups (p = 0.028), and on day 4 PI between the low and high groups (p = 0.042). Dashed lines represent mean percent weight, shaded regions indicate standard deviation, and open circles denote individual mice. **B) Disease Scores**: Higher *C. difficile* doses resulted in persistently elevated disease scores. Significant differences in clinical scores were observed on day 1 PI between the low and high groups (p = 0.026) and on day 4 PI between the low and medium (p = 0.011) and low and high groups (p = 0.008). Filled circles and error bars represent the median and interquartile ranges. Filled circles and error bars represent median and interquartile ranges. **C**) ***C. difficile* burden**: *C. difficile* burden initially correlates with inoculum size, but plateaus by day 4 PI. On day 1 PI, a significant difference in *C. difficile* burden was observed between the low and high (p = 0.004), and medium and high (p = 0.009) groups. The open circles represent individual mice.

To assess microbial composition, DNA was extracted from stool samples before and during CDI, and full-length 16S DNA was sequenced using long-read Oxford Nanopore sequencing. Several OMM19.1 strains were not detected in the stool samples (Figure 2). This is consistent with the original synthetic community construction by A. Afrizal et al. (2), who reported that *M. intestinale* was not detected after breeding and that *B. animalis* detected in OMM12 was not detected in OMM19.1 mice. The inability to detect *A. muris* may have been due to incomplete cell lysis.

**Figure 2.**
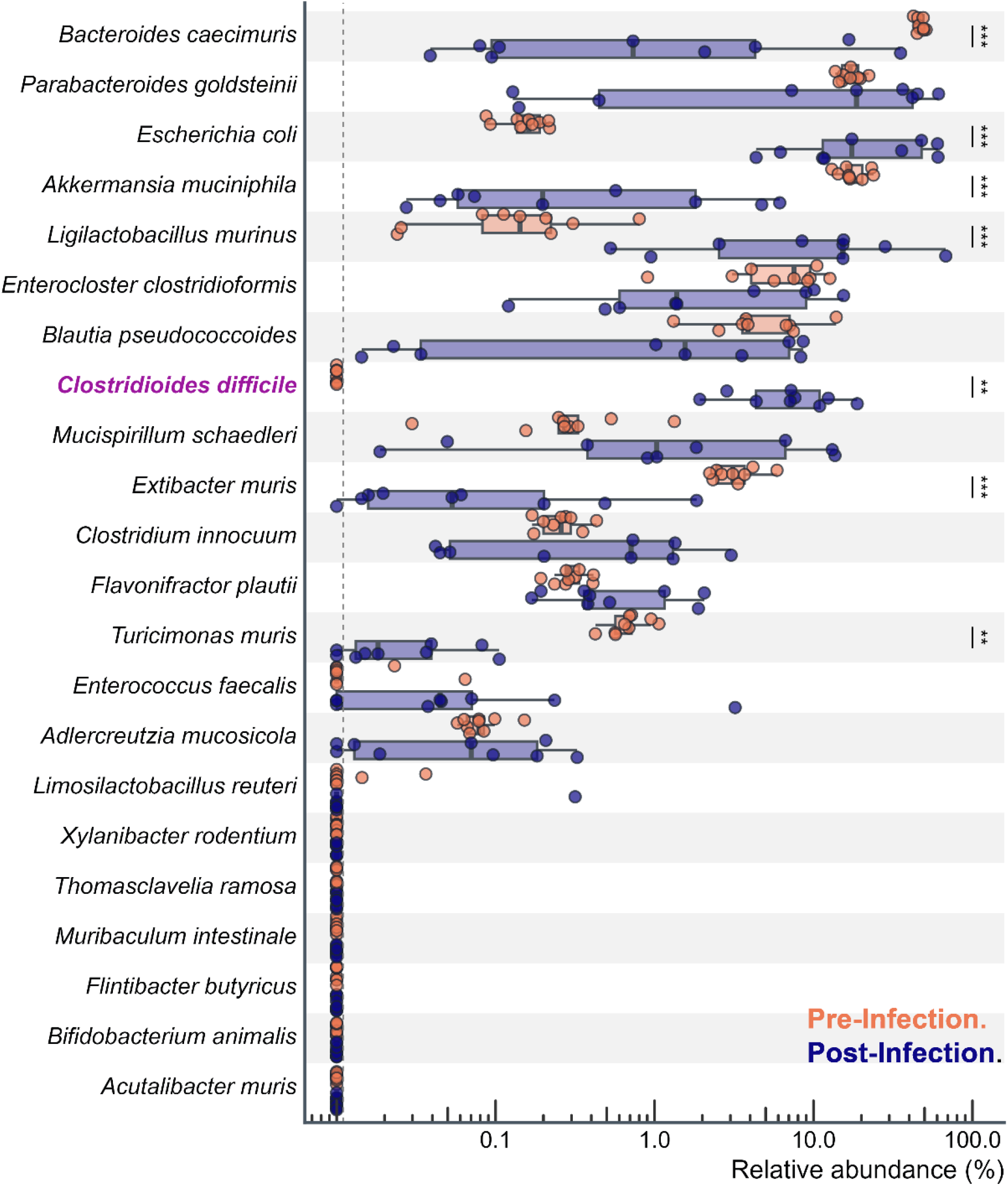
*C. difficile* infection significantly alters the OMM19.1 community without prior antibiotics. Infection with *C. difficile* resulted in a significant increase in *E. coli* and *L. murinus* while significant reductions in *B. caecimuris, A. muciniphila, E. muris*, and *T. muris* were observed. The dotted line indicates the limit of detection (e.g., no sequences were observed). The mice are represented by individual points (n = 22). p-value ≤ 0.01 = 2 stars (**). p-value ≤ 0.001 = three stars (***).

Previously, *A. muris* genomic DNA was isolated only when additional enzymatic steps were used (2). *T. ramosa* (*or Clostridium ramosum*) and *L. reuteri* were intermittently detected and observed at only one facility. *F. butyricus* was not detected in any of the OMM19.1 mice (2). Importantly, given that stool samples were sequenced, it is not possible to rule out the colonization of these strains further up the gastrointestinal tract or at low abundance not sampled by sequencing.

*C. difficile* infection significantly correlated with an increase in *Ligilactobacillus murinus* (log2 fold change (log2 FC) of 6.4 compared to uninfected, p < 0.001) and *Escherichia coli* (log2FC = 7.59, p < 0.001), and a decrease in *Bacteroides caecimuris* (log2FC = -2.83, p < 0.001), *Akkermansia muciniphila* (log2FC = -3.57, p < 0.001), *Extibacter muris* (log2FC = -3.51, p < 0.001), and *Turicimonas muris* (log2FC = -4.70, p = 0.004, Figure 2). While these changes are likely caused in part by an increase in intestinal inflammation, direct antagonistic or mutualistic interactions between *C. difficile* and the microbiota cannot be ruled out. Therefore, this model may provide a method to dissect these interactions. Interestingly, the supernatant from *L. murinus* culture has been demonstrated to significantly enhance the growth of *C. difficile* (6), while Proteobacteria are consistently increased in humans with CDI (7). Conversely, *Bacteroides, A. muciniphila*, and *E. muris* have been inversely associated with *C. difficile* and may ameliorate CDI (8-13).

## Conclusions

The OMM19.1 synthetic community builds upon OMM12 with additional strains covering a greater taxonomic range, including secondary bile acid producers and fiber degraders. The phenotype of OMM19.1 colonized mice has been shown to be more similar to SPF mice both physiologically, and immunologically. Colonization resistance is one of the strongest barriers to *C. difficile* infection, although the exact mechanisms remain to be determined. Despite being colonized by a functionally diverse community, OMM19.1 mice were susceptible to CDI. This may be attributed to several factors. Recent studies have suggested that the protective effects of *C. scindens* against *C. difficile* may be due to the production of the antimicrobial compound 1acetyl-β-carboline rather than inhibition by secondary bile acids (14). Furthermore, A. M. Aguirre et al. (15) showed that protection against CDI occurs without formation of secondary bile acids.

*C. difficile* is capable of Stickland metabolism where proline and glycine are reduced to 5-aminovalerate and acetyl phosphate, respectively. Thus, restoring the gut microbiome with bacteria that consume proline and/or glycine helps promote protection against CDI (15). The OMM19.1 community provides a good starting point for the elucidation of colonization resistance, but requires additional microbes.

### OMM19.1 Mice

The OMM19.1 community consists of 21 strains (Table 1) and was purchased, pre-mixed, from the DSMZ-German Collection of Microorganisms and Cell Cultures. Germ-free C57BL/6 mice were obtained from the Functional Microbiomics, Inflammation, & Pathogenicity Center (University of Louisville) and directly inoculated by gavage (50 μl orally, 50 μl rectally) with the OMM19.1 mixture (IACUC 21945). To ensure the complete transfer of microbes, inoculation was repeated 72 h after the initial inoculation. OMM19.1 mice were housed under gnotobiotic conditions and mated with the resultant F1 generation used for subsequent infection experiments. To confirm the colonization of the strains, fresh fecal pellets were obtained and frozen at −70 °C until needed.

### *C. difficile* infection and enumeration

F1 OMM19.1 mice of both sexes aged 6–8 weeks were challenged with a low (10^3^), medium (10^5^), or high (10^7^) dose of *C. difficile* 2015 (RT027) spores by oral gavage (IACUC 24362). *C. difficile* in stool was enumerated on *Clostridioides difficile* moxalactam norfloxacin agar supplemented with 0.1% sodium taurocholate (CDMNT) (16). The CDMNT plates were incubated anaerobically at 37 °C for 48 h prior to enumeration. Mice were weighed and monitored daily for signs of disease throughout the infection period. Clinical scores were determined using the criteria described by R. D. Shelby et al. (17).

### 16S gene sequencing

DNA was extracted from frozen fecal pellets using the DNeasy PowerLyzer PowerSoil Kit (Qiagen). DNA samples were sent to SeqCoast Genomics for sequencing using 16S full-length Nanopore reads (10k reads). Sequencing reads were analyzed with the Oxford Nanopore EPI2ME running the 16S workflow (wf-16s, v1.2.0) with the following settings: minimum reference coverage, 92%; minimum length, 1000 nt; maximum length, 1650 nt; and utilizing pysam (v0.21.0), pandas (v2.0.3), fastcat (v0.15.1), minimap2 (v2.26-r1175), samtools (v1.18), taxonkit (v0.15.1), and Kraken (v2.1.3).

### Statistical analysis

The non-parametric Kruskal–Wallis and Wilcoxon rank-sum tests were used for all statistical analyses, with Holm correction for multiple comparisons.

## ACKNOWLEDGMENTS

This work was supported by the Centers of Biomedical Research Excellence (CoBRE) Grant P20GM125504 and the Jewish Heritage Fund for Excellence Research Recruitment Grant Program at the University of Louisville, School of Medicine.

